# Stomatin encapsulates aquaporin-1 and urea transporter-B in the erythrocyte membrane

**DOI:** 10.1101/2025.08.29.673128

**Authors:** F. Vallese, H. Li, O.B. Clarke

**Affiliations:** Structural Biology Initiative, CUNY Advanced Science Research Center, New York, NY 10031, USA; Department of Chemistry and Biochemistry, City College of New York, New York, NY 10031, USA; Department of Anesthesiology, Columbia University Irving Medical Center, New York, NY,10032, USA; Department of Physiology and Cellular Biophysics, Columbia University, New York, NY, 10032, USA

## Abstract

Stomatin is a ubiquitous and highly expressed protein in erythrocytes, which associates with cholesterol-rich microdomains in the plasma membrane and is known to regulate the activity of multiple ion channels and transporters, but the structural basis of association with stomatin targets remains unknown. Here we describe high-resolution structures of multiple stomatin complexes with endogenous binding partners isolated from human erythrocyte membranes, revealing that stomatin specifically associates with two membrane proteins involved in water transport and cell volume regulation, aquaporin-1 (AQP-1) and the urea transporter, UT-B (SLC14A1). Together, our results reveal the structural basis of stomatin oligomerization, membrane association, and target recruitment, and identify a putative role for stomatin in the regulation of osmotic balance in the erythrocyte.

## Introduction

Compartmentalization is a fundamental principle of cell biology, enabling the spatial separation of distinct metabolic pathways and control of the local solution conditions. Apart from membrane-bound organelles, compartmentalization can be achieved by the assembly of protein microcompartments such as encapsulins in bacteria and archaea^1^, the soluble major vault protein in eukaryotes^2,3^, and a broad class of membrane-associated oligomeric assemblies including homologs of Stomatin, Prohibitin, Flotillin, and HflK/C, known as the SPFH superfamily. The first and eponymous member of the SPFH family, stomatin (STOM) was first identified in erythrocytes (where it is also known as band 7), and is expressed in many cell types and organisms^4^. Stomatin deficiency has been linked to erythrocyte fragility and a condition called overhydrated hereditary stomatocytosis, in which erythrocytes leak Na^+^ and K^+^ ions ^5^. Since the identification of stomatin, numerous SPFH family members have been identified in all three domains of life. The conservation among these proteins is high, with human stomatin homologs sharing >50 % identity^6^. Despite being associated with a variety of diseases such as cancer, kidney failure, and anemia^7,8^, the molecular functions of SPFH family members in most cases remains enigmatic.

SPFH proteins regulate various cellular processes across all domains of life, from bacteria to eukaryotes^9^. Members of this family typically possess a single transmembrane or intramembrane (TM/IM) segment that anchors the protein to the membrane, as well as a conserved pair of cytosolic SPFH1/2 domains that mediate, along with a pair of coiled-coil domains, the formation of pseudosymmetric dome-like assemblies^10,11^. Bacterial SPFH proteins such as HflK/C are known to encapsulate target membrane proteins (such as the FtsH protease in the case of HflK/C) as one aspect of their mechanism for regulating target protein function, but the extent to which this mechanism applies to eukaryotic SPFH proteins is unclear. Cryo-EM structures of several eukaryotic SPFH proteins, including an apo-structure of recombinant stomatin^12^, were recently published^13^, showing that while the dome architecture and domain arrangement is largely conserved, the size, stoichiometry and interprotomer interfaces of the complex vary substantially between different family members.

Stomatin localizes to lipid rafts in the plasma membrane^14^ and is enriched in extracellular vesicles, consistent with a role in membrane organization and cargo sorting^15,16^. Functionally, stomatin and stomatin-like proteins have been implicated in channel/transporter modulation and mechanosensory pathways^16,17^. Reported interactors include Aquaporin-1 (AQP1), anion and cation channels (e.g., ASICs), pannexins, and glucose transporters ^18,19^, among others, though the molecular basis for these interactions has remained unclear.

Using single-particle cryo-EM, we have determined the architecture of stomatin complexes purified from native human erythrocyte membranes at an overall resolution of 2.0 Å. The stomatin oligomer forms a 16-subunit ring built from eight asymmetric homodimers, with C16/C8 mixed symmetry. We identify and characterize complexes with two endogenous partners, positioned beneath the dome, by using focused 3D classification: an AQP1 tetramer and a UT-B trimer surrounded by an encircling lipid belt. These data show that endogenous stomatin acts as a membrane scaffold that corrals and positions stomatin-associated membrane proteins, providing a structural framework for membrane protein microcompartmentalization at the erythrocyte plasma membrane.

## Results

### Purification and structure determination of human stomatin complexes

Stomatin complexes were purified from digitonin-solubilized human erythrocyte membranes by the following strategy. First, high molecular weight complexes (including the ankyrin complex and the stomatin complexes) were separated from lower molecular weight species by glycerol density gradient centrifugation. The stomatin complexes were then separated from other contaminants by anion exchange, followed by size exclusion chromatography, to generate the final sample used for structural characterization (**Extended Data** Fig. 1). Initial structural characterization of the digitonin-solubilized stomatin complex revealed an apparently C16 symmetric, dome-shaped assembly, with a partially membrane-embedded N-terminal SPFH1 domain (**Extended Data** Fig. 3 **a**). Featureless beta strands on the inner wall of the apical cap region led us to suspect the presence of pseudosymmetry, and focused classification without alignments allowed us to identify the overall symmetry as C8 (**Extended Data** Fig. 2), with 8 asymmetric homodimers forming the 16-protomer assembly. Refinement in C8 resulted in a reconstruction with an overall resolution of 2.0 Å, and facilitated building a near complete model of human stomatin (**Fig. 1**), including multiple palmitoylation sites in the membrane-embedded region.

**Fig. 1:**
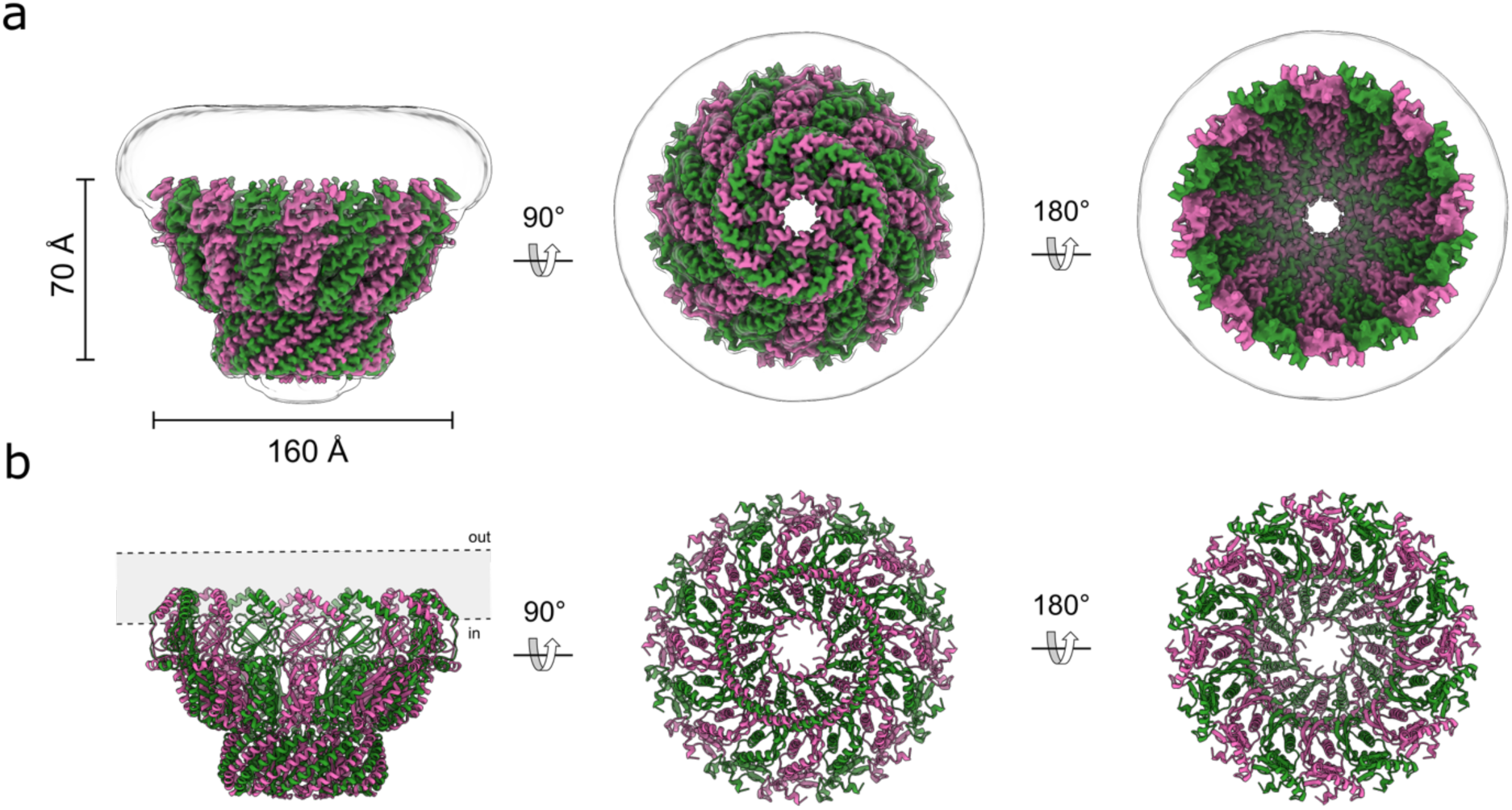
Architecture of human stomatin. **a.** Cryo-EM density map of stomatin viewed in the membrane plane (left), from the cytoplasmic side (center), and the extracellular side (right). The 16 stomatin protomers are alternately colored pink (protomer A) and green (protomer B). The Gaussian-filtered map is shown as a transparent surface to delineate the boundaries of the micelle. **b,** Atomic model of stomatin, shown in the same orientations and color scheme as in panel a.

### Architecture and pseudosymmetry of the stomatin dome

The cryo-EM reconstruction reveals a dome-shaped 16-subunit ring assembled from eight asymmetric homodimers, with C16/C8 mixed symmetry (**Fig. 2 a**). The membrane-proximal base is formed by an SPFH1 domain and an intramembrane helix (IMH1; residues ∼20–53) contributed by each protomer, which together define the base of the dome (**Extended Data** Fig. 3 **a**). The cytosolic SPFH2 domains form a second layer of interactions at the base of the dome, followed by the first coiled coil (CC1) domains which arch upward to an apical cap (distal to the membrane), formed by the second coiled coil domain (CC2) and the C-terminal beta strand (beta-Ct). The beta-Ct strand forms a 16-stranded beta barrel, in which every adjacent strand (contributed alternately by the A and B protomers) is offset in sequence register with respect to its neighbor by one residue (**Fig. 2 b**). At 2.0 Å overall resolution, side chains and solvent molecules are well resolved throughout the dome and intramembrane belt (**Extended Data** Fig 3 **d & Movie S1-S2**). The only poorly ordered region is the very N-terminus, where lower-contour views reveal weak density for the first ∼20 residues which bind to the SPFH1/2 interdomain interface. The base spans ∼140 Å outer diameter; the SPFH belt widens to ∼160 Å before tapering toward the apex, yielding a dome height of ∼100 Å. A continuous axial pore runs through the assembly and narrows to a ∼7.3 Å final radius in the C-terminal beta barrel (**Extended Data** Fig. 3c), which is lined almost exclusively by hydrophobic residues (**Fig. 2b & Extended Data** Fig. 3 **b**). The top pore remains open, showing only faint density and a few unmodeled C-terminal residues, with no ordered plug, suggesting the possibility that this aperture may be permeable to solvent (**Extended Data** Fig. 3 **e**). Contiguous density is observed protruding from the sidechains of Cys30, Cys53, and Cys87, compatible with S-palmitoylation (**Fig. 2c & Extended Data** Fig. 10). This is consistent with prior literature for Cys30 and Cys87^20,21^, and we now see analogous density at Cys53, suggesting possible palmitoylation of this residue also. Together, these modifications could help anchor the complex in lipid rafts and promote stability of the dome. A local asymmetric refinement of the membrane-embedded region also allows us to identify multiple non-covalently bound phospholipids, one of which is consistent with sphingosine-1-phosphate (**Fig. 2d**). Perhaps surprisingly given the association of stomatin with cholesterol rich domains in the membrane, no ordered cholesterol molecules are identified binding to the stomatin intramembrane region. In the C-terminal cytosolic beta barrel, eight non-protein densities are found lining the hydrophobic aperture. Their shape is inconsistent with cholesterol or digitonin and appears more consistent with a free fatty acid, although they cannot be unambiguously identified (**Extended Data** Fig. 3 **e**).

**Fig. 2:**
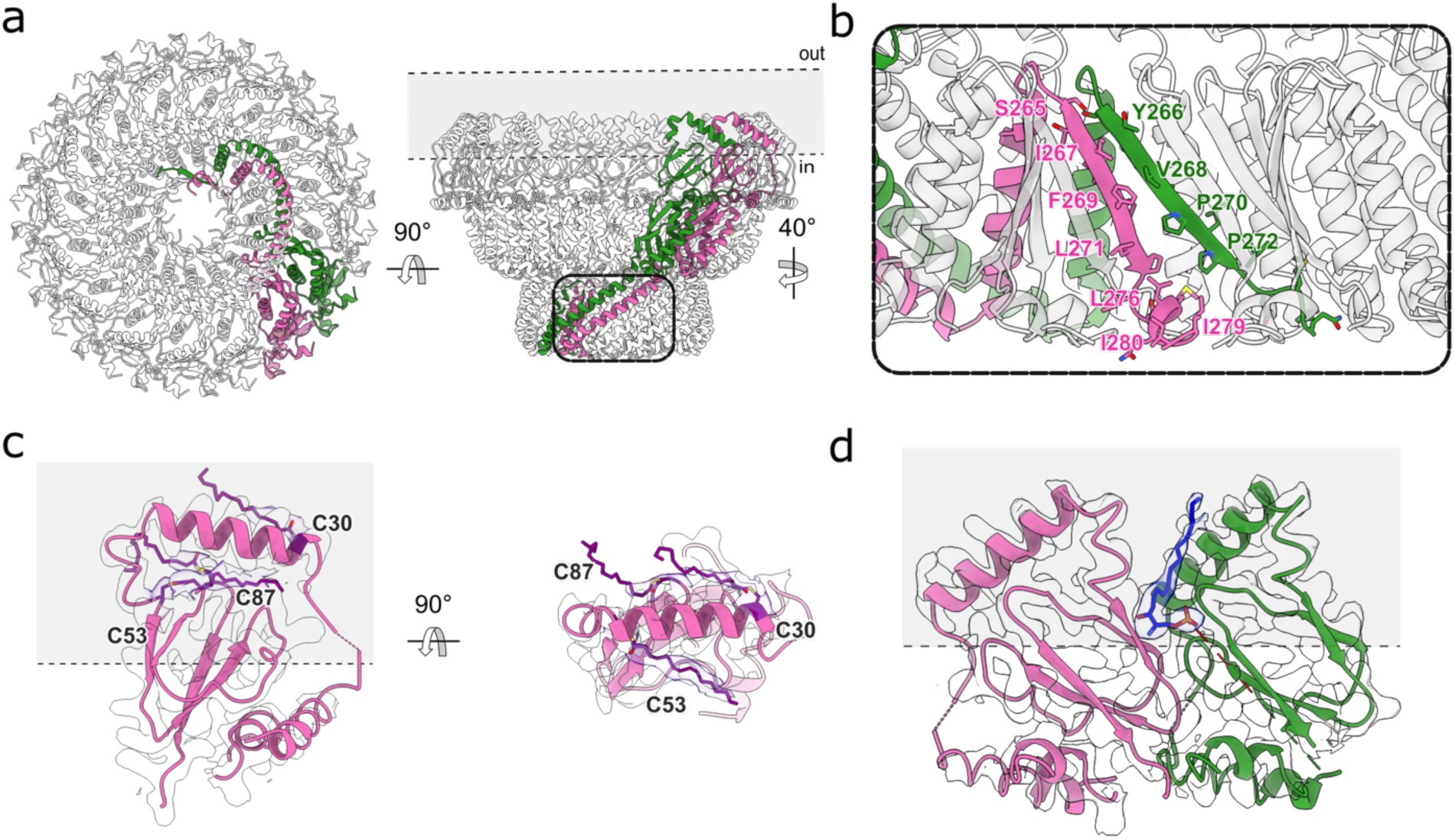
Structural features of stomatin. **a** Model of stomatin with one of the eight dimers highlighted (pink and green for stomatin protomers A & B respectively), viewed from the cytoplasmic side (left) and in the membrane plane (right). **b,** Close-up view of the stomatin apical C-terminal beta-barrel, showing residues that line the internal pore. **c,** Atomic model of the SPFH1 domain and IMH1/2 region of stomatin protomer A viewed in the membrane plane, with putative palmitoylation sites on Cys30, Cys53, and Cys87 highlighted in purple, and the cryo-EM density map shown as a transparent surface. **d**, Model of the intramembrane region of a dimer, with the sphingosine-1-phosphate modeled between the monomers shown in blue.

### Identification of endogenous stomatin binding partners by deep 3D classification

Filtering the C8 or C16 reconstructions to low resolution revealed strong, but uninterpretable density in the center of the digitonin micelle, leading us to suspect the presence of stomatin binding partners under the dome. Classification of asymmetric transmembrane binding partners associating with a highly symmetric, but flexible assembly is a challenging image processing task, and initial classification attempts using multiple different approaches failed or gave ambiguous results. Focused classification without alignments using an unusually large number of 3D classes (which we are terming here deep 3D classification) at low resolution (12-15Å) succeeded in identifying two main populations of stomatin-encapsulated membrane proteins – one with apparent local C4 symmetry, and another with apparent C3 symmetry (**Extended Data** Fig. 4 **& 5**). Local refinement with the apparent local symmetry enforced facilitated high resolution reconstructions of both species, identifying the former as aquaporin-1 (AQP1), resolved at 2.3Å, and the latter as the urea transporter UT-B, resolved at 2.8Å (**Fig. 3**). Based on analysis of the classification results, approximately 40% of stomatin domes are associated with AQP1, 10% are associated with UT-B, and 50% are empty or have inconclusive density. Aquaporin-1 is positioned asymmetrically within the stomatin dome, interacting directly with the inner portion of the intramembrane domain (**Extended Data** Fig. 6 **a, b**), while the UT-B trimer is positioned in the center of the stomatin dome, interacting with stomatin via lipid-mediated interactions (**Extended Data** Fig. 6 **c**).

**Fig. 3:**
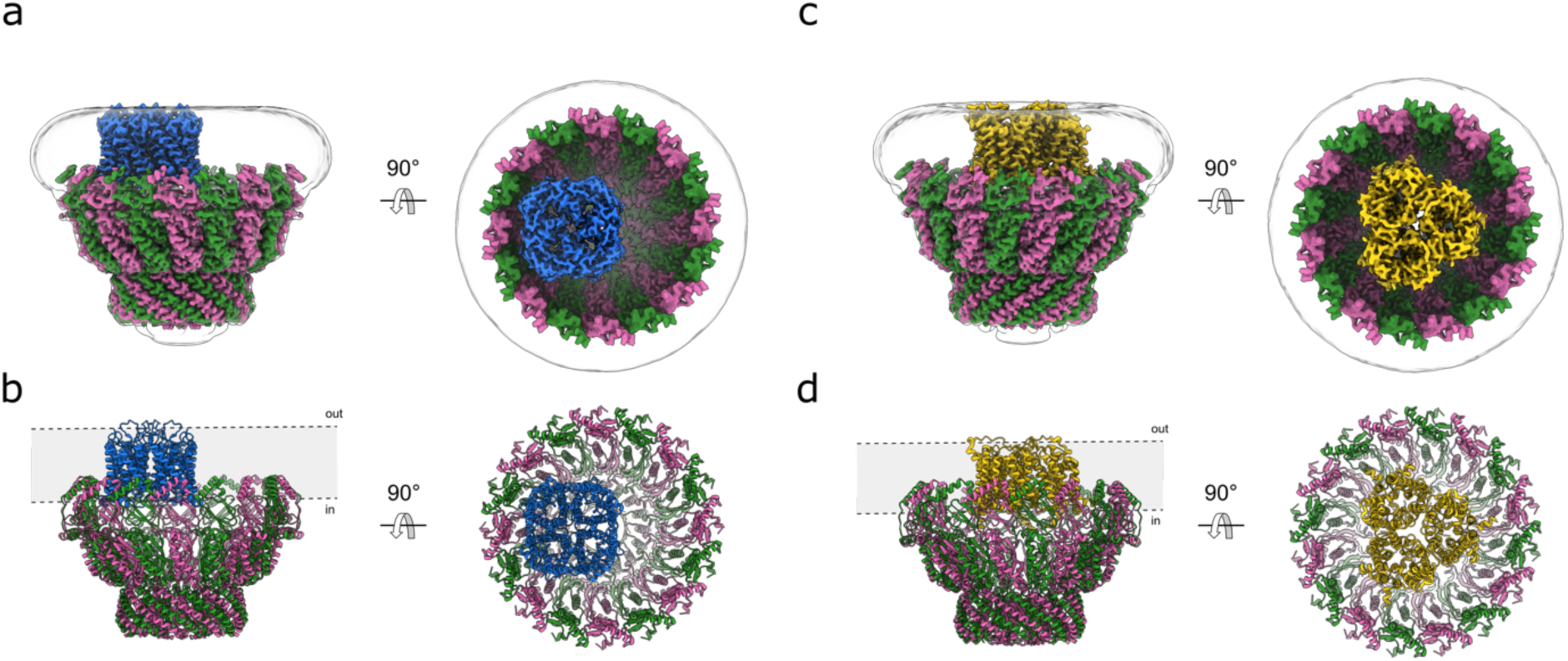
Structural identification of AQP1 and UT-B as stomatin associated membrane proteins. **a,** Cryo-EM density map of stomatin (pink and green) in complex with AQP1 (blue), viewed in the membrane plane (left) and from the extracellular side (right). The Gaussian-filtered map delineating the micelle boundary is shown as a transparent white surface. **b,** Atomic model of the stomatin-AQP1 complex, shown in the same orientations and color scheme as in panel a. **c,** Cryo-EM density map of stomatin (pink and green) in complex with UT-B (yellow), viewed in the membrane plane (left) and from the extracellular side (right). **d,** Atomic model of the stomatin-UT-B complex, shown in the same orientations and color scheme as in panel a.

### Architecture of the AQP1 & UT-B stomatin complexes

A single AQP1 tetramer is located within the membrane patch defined by the stomatin dome, interacting with the wall of the dome, primarily with IMH2 (**Fig. 3 a,b & Extended Data** Fig. 6 **a,b**). No classes were identified with multiple AQP1 tetramers. Each AQP1 monomer contains a single water pore, so the tetramer harbors four independent water pathways. As identified in the structure of ankyrin-bound AQP1^22^, Cys87 of AQP1 is palmitoylated, and a bound cholesterol molecule is observed near the intracellular interprotomer interface. In the stomatin-AQP1 structure, the tetramer sits off-axis relative to the assembly’s pseudo-C16 symmetry axis and engages the intramembrane surfaces of two stomatin protomers belonging to adjacent dimers (**Extended Data** Fig. 6 **b**). The stomatin intramembrane belt forms a continuous ring that surrounds AQP1, effectively sequestering it from the surrounding bilayer and limiting lateral diffusion outside of the lumen. The protein–protein interface is limited, probably confined to AQP1 cytosolic loops and termini (**Extended Data** Fig. 6 **a,b**); the water pore remains fully resolved and unobstructed in each monomer.

3D classification also revealed a second partner within the dome: the urea transporter UT-B (SLC14A1) (**Fig. 3 c,d**). In contrast to AQP1, the homotrimeric UT-B transporter is positioned in the center of the dome, with each subunit contributing 10 transmembrane helices and the trimer presenting its characteristic triangular footprint to the membrane. We observe a continuous annulus of density surrounding the UT-B transmembrane surface (**Extended Data** Fig. 6 **c**). Although we cannot unequivocally assign the outer lipids, they form a “lipid belt” that seems to bridge UT-B to the inner face of the stomatin intramembrane ring, providing lipid-mediated contacts in the membrane plane. We do not resolve direct polypeptide–polypeptide contacts between UT-B and stomatin within the bilayer, with all contacts appearing to be lipid-mediated. The geometry and location of many of the phospholipids that directly contact UT-B match those reported in recent human UT-B/UT-A structures, where lipids occupy defined grooves between subunits^23^.

## Discussion

Our structures define stomatin as a dome-shaped, C16/C8 pseudosymmetric assembly that corrals specific membrane proteins within a membrane patch under the stomatin dome. Two native stomatin associated membrane proteins, AQP1 and UT-B, sit inside the intramembrane belt beneath the dome, providing a direct structural basis for stomatin-dependent regulation of water and urea transport in the erythrocyte membrane, processes critical for regulation of osmotic balance and cell volume ^24,25^. Beyond erythrocytes, these data support a general model in which SPFH oligomers build membrane-bound microcompartments that couple tuning of local lipid composition to cargo selection and function. An analogous “membrane-encapsulin” role has been characterized in bacteria, where the SPFH pair HflK/C assemble a ring that corrals multiple FtsH proteases into a microcompartment, spatially organizing proteolysis at the membrane^26^.

SPFH proteins all possess a single intramembrane or transmembrane segment and a conserved pair of SPFH1 and SPFH2 domains, yet their higher-order architectures are diverse (**Extended Data** Fig. 7). Stomatin forms a ∼160 Å dome from eight asymmetric homodimers, with a narrow, hydrophobic apical pore and a continuous intramembrane belt. Differences in ring diameter, pore geometry, and belt topology (which can be transmembrane or intramembrane), as well as the lipid composition of the host membrane, likely tune the cargo selectivity of different SPFH family members.

Deep 3D classification of native stomatin, purified from human erythrocytes, resolved two stomatin-cargo complexes, with AQP1 and UT-B, demonstrating that stomatin accommodates cargos of distinct symmetry and interaction modes, which may be direct (AQP1) or lipid-mediated (UT-B). Characterization of the stomatin-AQP1 and stomatin-UT-B complexes extends the encapsulin principle to the eukaryotic plasma membrane, with the stomatin oligomer forming a palmitoylated fence that isolates cargos in raft-like regions and dampens their lateral mobility. The deep 3D classification approach we have applied here to stomatin, may be useful for resolving the cargo of other native SPFH complexes for which data is available, such as the flotillin1/2 complex. It is unclear whether encapsulation is the only mechanism for stomatin regulation, or whether other target proteins may associate with the outside of the dome. The hydrophobic apical pore, which we see here is decorated by small amphiphiles, likely forms a conduit for the transport substrates of the resident membrane proteins, although it remains possible, depending on the conformation of the unresolved C-termini, that the cap may seal off the cargo membrane protein from the surrounding solvent, either temporarily/dynamically or permanently (**Fig. 4**).

**Fig. 4:**
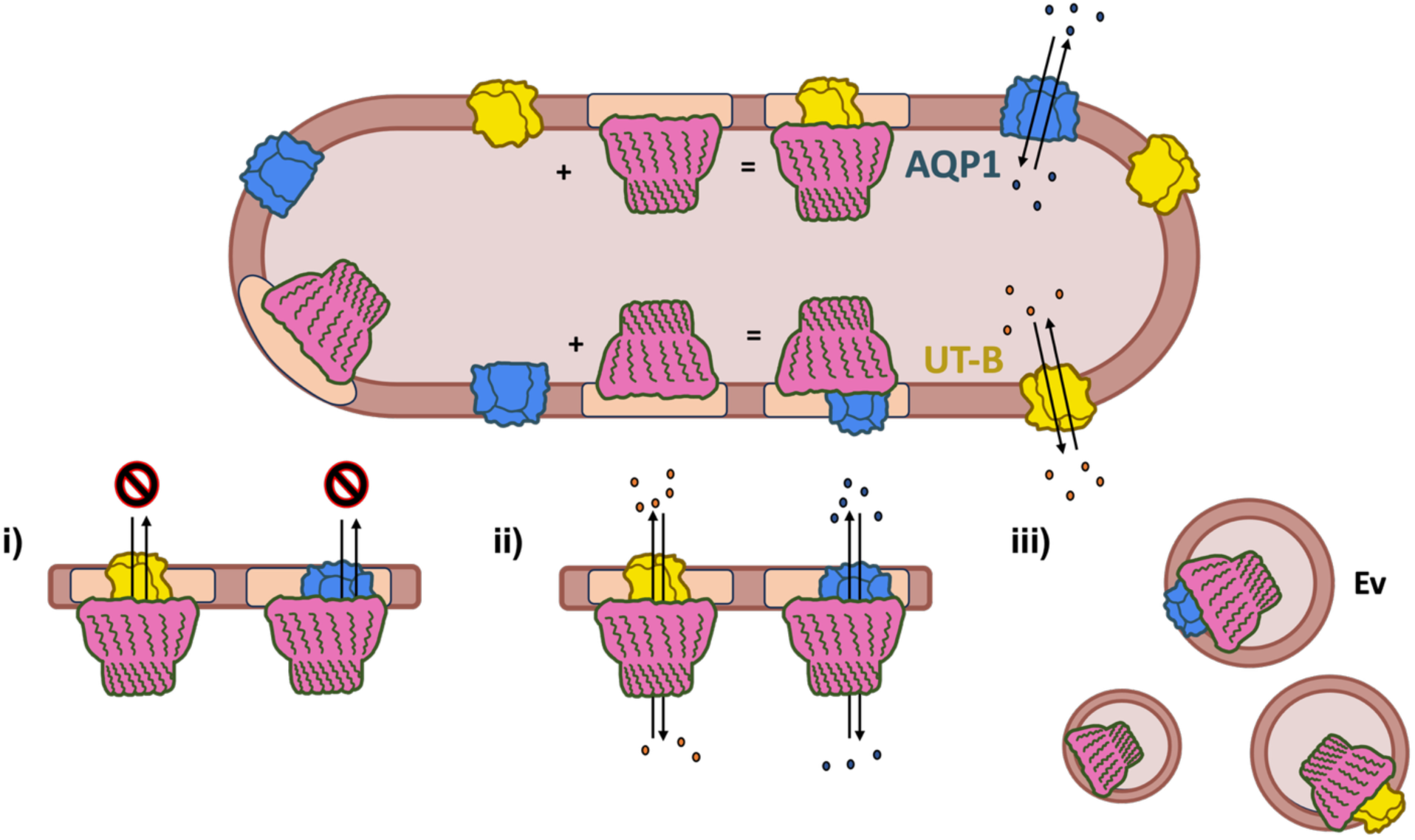
Schematic of the human stomatin in erythrocytes. Illustration of the possible functions of stomatin complex formation with AQP1 (blue) and UT-B (yellow) in erythrocytes. Beige shading in the membrane indicates microdomains. Stomatin may regulate AQP1 and UT-B in several ways: (i) blocking substrate exchange, water for AQP1 and urea for UT-B; (ii) capping the proteins to modulate substrate flux by restricting flow of substrates into and out of the stomatin dome; and (iii) promoting removal of cargo proteins from the plasma membrane via extracellular vesiculation.

While the structural basis for stomatin association with target membrane proteins, and the identity of two of them, is now clear, several questions remain to be answered regarding the mechanistic basis and functional consequences of this process. Firstly, questions surround the timing and dynamics of stomatin complex assembly. Erythrocytes do not have ribosomes (or if they do, they exist at a very low level for specialized purposes^27^), so when are the stomatin domes being assembled around target proteins? If it is happening in the mature erythrocyte, this implies the existence of a pool of partially assembled or disassembled stomatin domes, either in the cytosol or at the membrane, which can facilitate the entry of AQP1 or UT-B. Alternatively, it is possible that stomatin-encapsulated membrane proteins are assembled co-translationally, during erythrocyte biogenesis, and perhaps the stomatin dome is then degraded by erythrocytic proteasomes^28^ during erythrocyte aging^29^, releasing target membrane proteins. In terms of the functional consequences of stomatin association, some clues exist, although no definitive answers are yet available. Firstly, during overhydrated hereditary stomatocytosis, which is caused by mutations not in stomatin, but in the heterotrimeric ammonia transporter RhAG^30^, stomatin is initially present, in young erythrocytes, at near-normal levels; but it is progressively lost during erythrocyte aging^31^. Secondly, high levels of stomatin, along with stomatin associated membrane proteins, are known to be present in extracellular vesicles which are secreted from erythrocytes during aging or blood storage^32^. These two observations taken together suggest a possible role for stomatin in sequestration and/or disposal of unneeded or damaged targeted membrane proteins.

Erythrocytes experience extreme mechanical stress and rapid osmotic shifts yet lack endomembranes and transcriptional plasticity, necessitating alternative methods to control membrane transport processes. Stomatin provides (i) compartmentalized control of water and urea flux via AQP1 and UT-B, (ii) possible stabilization of raft architecture, and (iii) a sorting platform that can be shed in extracellular vesicles during blood aging or storage (**Fig. 4**), together offering a plausible basis for coordinated water/urea handling in circulation.

## Acknowledgments

Cryo-EM data were collected at the Columbia Cryo-EM facility and at the Simons Electron Microscopy Center (SEMC), with the assistance of staff from both SEMC and the Columbia University Cryo-Electron Microscopy Center. R. Grassucci and Z. Zhang from the Columbia Cryo-EM Center assisted with data collection. This study was supported by the National Institutes of Health (NIH) through grants R01-HL168178 (O.B.C). The Irma T. Hirschl Trust provided support for O.B.C. We thank Kookjoo Kim for the support. We thank Tito Calì (U. Padova) for helpful discussions and advice.

## Author Contribution

F.V. and O.B.C. conceived the study. F.V. and O.B.C. designed the experiments. F.V. and O.B.C. performed structural biology experiments, and O.B.C. built and refined structural models. Figures were prepared by F.V. and H.L. All authors analyzed the results. F.V. and O.B.C. wrote the manuscript.

## Methods

### Stomatin sample preparation for cryo-EM

The general purification workflow is presented in Extended Data Fig. 1. Erythrocyte ghost membranes were prepared as described by Niggli et al. 2016^33^. Briefly, red blood cells from healthy blood donors were purchased from the New York Blood Center. Erythrocytes were washed twice in 5 volumes of 130 mM KCl, 10 mM Tris-HCl, pH 7.4. The cells were hemolyzed in 5 volumes of 1 mM EDTA, 10 mM Tris-HCl, pH 7.4, and centrifuged at 18,000 × g for 10 min. The resulting ghost membranes were then washed five times in the hemolysis buffer and four additional times in 10 mM HEPES, pH 7.4. The hemoglobin-free ghost membranes were finally resuspended in 130 mM NaCl, 20mM HEPES, pH 7.4, 0.5 mM MgCl_2_, 0.05 mM CaCl_2_, 2 mM dithiothreitol (DTT), and stored at –80°C.

Ghost membranes were solubilized at a protein concentration of 4 mg/ml in 130 mM KCl, 10 mM HEPES, pH 7.4, protease inhibitor tablet (cOmplete^™^, EDTA-free Protease Inhibitor Cocktail, Millipore Sigma), 1% (w/v) digitonin (Carbosynth), at 4°C for 1 hour. Unsolubilized material was removed by centrifugation at 26.000 × g for 30 min. The supernatant was applied to a PD10 column (to reduce the detergent concentration) equilibrated with 0.05% (w/v) digitonin, 130 mM KCI, 20 mM HEPES, pH 7.4. The sample was than applied on a glycerol step gradient (30– 12% glycerol) and centrifuged for 15 h at 25,000 rpm (SW 32 rotor, Beckman). The distribution of stomatin was confirmed by SDS-PAGE gel (4–20% Mini-PROTEAN TGX Precast Protein Gels, Bio-Rad). Fractions containing stomatin were pooled together and concentrated to <500 μL using a 100-kDa cutoff concentrator. To further purify the sample, it was loaded onto a Superose 6 10/300 Increase column (Cytiva) equilibrated in 0.05% (w/v) digitonin, 130 mM KCl, and 20 mM HEPES, pH 7.4. Fractions enriched in stomatin were pooled and concentrated to 1 ml. The sample was then applied to a PD10 column equilibrated with 0.05% (w/v) digitonin and 20 mM Tris-HCl, pH 6.5, and subsequently loaded onto a 5 mL HiTrap ANX FF column (GE Healthcare). Proteins were eluted from the column with a step gradient of 0–0.6M KCl in the same buffer (0.15M, 0.3M and 0.6M). The fractions from the 0.3M KCl elution containing stomatin were collected and concentrated to ∼<500 μL using a 100-kDa cutoff concentrator (Millipore). The concentrated sample was further subjected to size-exclusion chromatography on a Superose 6™ 10/300 column (GE HealthCare) equilibrated with 0.05% (w/v) digitonin, 130 mM KCI, 20 mM HEPES, pH 7. The fraction enriched in stomatin was immediately used for cryo-EM grid preparation.

### Cryo-EM grid preparation and data collection

3 µL of purified stomatin at 4 mg/mL with a surface-active additive (0.01% (w/v) glycyrrhizic acid) was added to a glow discharged (PELCO easiGlow) 0.6/1 µm holey gold grid (Quantifoil UltrAuFoil) and blotted for 4-6 s at 4°C and 100% humidity using the Vitrobot Mark IV system (ThermoFisher Scientific), before plunging immediately into liquid ethane for vitrification. The cryo-EM data were collected on a Titan Krios electron microscope (ThermoFisher Scientific) equipped with a K3 direct electron detector (Gatan) operating at 0.83 Å per pixel in counting mode using Leginon automated data collection software^34,35^. Data collection was performed using a dose of 58 *e*^−^/Å^2^ across 50 frames (50 ms per frame) at a dose rate of 16 *e*^−^/pixel/s, using a set defocus range of –0.5 to –1.5 µm. A 100-µm objective aperture was used. A total of 20,647 micrographs were collected.

### Cryo-EM data processing

#### Pre-processing, initial volume generation and C16 consensus refinement

The final cryo-EM data processing workflow is summarized in (**Extended Data** Fig. 2). Orientation distributions, FSC plots, and validation statistics are presented in **Extended Data** Fig. 8 **& 9** and **Table 1**. Maps, masks and raw movies have been deposited at EMDB (IDs: TBD) and EMPIAR (IDs: TBD). Subsequent steps were performed in cryoSPARC v4.4-4.71 unless otherwise indicated^36^.

**Table 1.** Cryo-EM data collection, refinement, and validation statistics.

|  | **Stomatin** | **AQP1** | **UT-B** | **AQP1** | **Stomatin** |
| --- | --- | --- | --- | --- | --- |
|  | C8 | C4 | C3 | C1 | **intramembrane** |
|  | PDB | PDB | PDB | PDB | PDB |
|  | EMD- | EMD- | EMD- | EMD- | EMD- |
| **Data collection and processing** |  |  |  |  |  |
| Magnification | 105,000x | 105,000x | 105,000x | 105,000x | 105,000x |
| Voltage (kV) | 300 | 300 | 300 | 300 | 300 |
| Electron exposure (e<sup>-</sup>/Å<sup>2</sup>) | 58 | 58 | 58 | 58 | 58 |
| Defocus range (μm) | -0.5 to -1.5 | -0.5 to -1.5 | -0.5 to -1.5 | -0.5 to -1.5 | -0.5 to -1.5 |
| Pixel size (Å) | 0.83 | 0.83 | 0.83 | 0.83 | 0.83 |
| Symmetry imposed | C8 | C4 | C3 | C1 | C1 |
| Initial micrographs | 20,647 | 20,647 | 20,647 | 20,647 | 20,647 |
| Final micrographs | 14,499 | 14,499 | 14,499 | 14,499 | 14,499 |
| Initial particle images | 592k | 592k | 592k | 592k | 592k |
| Final particle images | 38k | 135k | 29k | 110k | 135k |
| Subparticle images | N/A | 631k | 35k | 135k | 2.2M |
| Map resolution (Å) | 2 | 2.3 | 2.8 | 3.1 | 2.2 |
| **Model composition** |  |  |  |  |  |
| Non-hydrogen atoms | 33,848 | 7,464 | 8,355 | 14,799 | 6,340 |
| Protein residues | 4,368 | 988 | 979 | 2,933 | 780 |
| **RMS Deviations** |  |  |  |  |  |
| Bond lengths (Å) | 0.2 | 0.16 | 0.13 | 0.6 | 0.18 |
| Bond angles (°) | 0.42 | 0.32 | 0.31 | 0.92 | 0.36 |
| **Validation** |  |  |  |  |  |
| Clashscore | 6 | 4 | 7 | 9 | 6 |
| Poor rotamers (%) | 1 | 1 | 1 | 1 | 1 |
| **Ramachandran plot** |  |  |  |  |  |
| Favored (%) | 96 | 97 | 98 | 96 | 94 |
| Allowed (%) | 4 | 3 | 2 | 4 | 6 |
| Disallowed (%) | 0 | 0 | 0 | 0 | 0 |

Patch-based motion correction and dose-weighting of 20,647 movies were carried out in cryoSPARC using the Patch Motion job type. Patch-based CTF estimation was performed on the aligned, non-dose-weighted averages using Patch CTF. Particles were manually picked from 93 micrographs. Topaz was run in training mode using a downsampling factor of 10 and an estimated number of particles per micrograph of 50. The resulting model was used to pick particles from the entire dataset using Topaz Extract. 592k particles were initially extracted in a box of 512 pixels and Fourier cropped to 96 pixels. Multiple rounds of 2D classification were performed to isolate homogeneous subsets of particles to use for ab initio reconstruction. Heterogeneous ab initio reconstruction was performed on 355k particles selected by 2D classification, first without symmetry and then, after testing multiple different Cn symmetry groups, with C16 symmetry, which gave a map with clear secondary structural features. An initial refinement was performed using non-uniform refinement^37^ (with C16 symmetry enforced) with on-the-fly refinement of per-particle defocus, and global refinement of beam tilt and trefoil aberrations, resulting in an initial consensus refinement with a resolution of 1.9 Å. Reference based motion correction improved resolution in C16 to 1.84 Å.

#### C8 symmetry determination using 3D Classification

In the C16 consensus non-uniform (NU) refinement map, the interior of the apex of the stomatin cap appeared poorly resolved, with no sidechain detail, while the immediately surrounding regions exhibited excellent density quality, leading us to suspect the presence of unresolved pseudosymmetry. To resolve the suspected pseudosymmetry in this region, a local mask covering this region was generated using UCSF ChimeraX^38–40^ and CryoSPARC^36^. Focused 3D classification on the cap region without alignments in C1 revealed two C8-symmetric classes. The classes were aligned using Align 3D maps in CryoSPARC, updating particle alignments accordingly, and subjected to local refinement using a global mask, with C8 symmetry enforced, resulting in a reconstruction with a resolution of 1.9 Å. The membrane embedded and membrane proximal regions of stomatin remained less well resolved than the rest of the oligomer, likely due to continuous deformation of the stomatin cap, as assessed by both 3D Variability Analysis and 3D classification with a ring-shaped mask covering the membrane-embedded region of stomatin (**Extended Data** Fig. 2). In order to obtain a map that allowed confident model building in the membrane embedded portion, another round of 3D classification with a mask around the stomatin membrane embedded region allowed the identification of a set of 38k particles with improved density for the intramembrane region, resulting in a 2.0 Å map after refinement with C8 symmetry enforced, with well-defined density for the intramembrane helix, palmitoylation sites, and bound lipids.

#### Identification of AQP-1 and UT-B using 3D Classification

To identify putative stomatin-associated membrane proteins, we applied 3D classification with 80 classes on a C16 symmetry expanded particle set using a featureless, disk-shaped mask centered on the membrane-embedded region, with a target resolution of 12 Å (**Extended Data** Fig. 4). Different numbers of classes were tested, and a number of classes greater than 40 was found to be necessary to separate the bound membrane proteins. This analysis revealed two main binding partners present in distinct complexes: tetrameric AQP-1 and trimeric UT-B. An overview of all 80 classes is shown in **Extended Data** Fig. 5. After aligning the rotationally-related classes and modifying the particle orientations accordingly, local refinement was employed in two stages to improve the density for AQP1 and UT-B, initially in C1, and then by applying the local molecular symmetry - C4 symmetry for AQP-1 and C3 symmetry for UT-B, resulting in maps at 2.3 Å and 2.8 Å resolution, respectively. To resolve the stomatin–AQP1 interface, we performed local asymmetric refinement starting from the initial post-classification reconstruction. A soft mask was generated to include the two AQP1 monomers in direct contact with stomatin, together with the four adjacent stomatin intramembrane domains. Local refinement was performed without symmetry enforced to improve the density at the AQP1-stomatin interface, resulting in a 3.1 Å map with significantly improved density for the contact region.

#### Atomic model building and refinement

An initial model for the stomatin protomer was generated using AlphaFold 2^41^. An AQP1 structure^22^ (PDB 7UZE) and UT-B structure^23^ (PDB 8XDF) were used as initial models for building the corresponding proteins. Initial models were placed in corresponding local reconstructions and fit as rigid bodies using `fitmap` in UCSF Chimera^42^. Each model was then manually extended and completed in COOT^43^. Residues 5-22 of stomatin were assigned as UNK residues to indicate that the sequence register in this region is ambiguous. Waters and ligands were placed where justified by the density and chemical environment, and each final model was refined against the respective map using `phenix.real_space_refine` ^44^. An overview of the model-density fit for the different structures is provided in **Extended Data** Fig. 10. Figures were prepared using UCSF ChimeraX^38–40^. Refinement and validation statistics are provided in **Table 1**.

---

**Extended Data Fig. 1:**
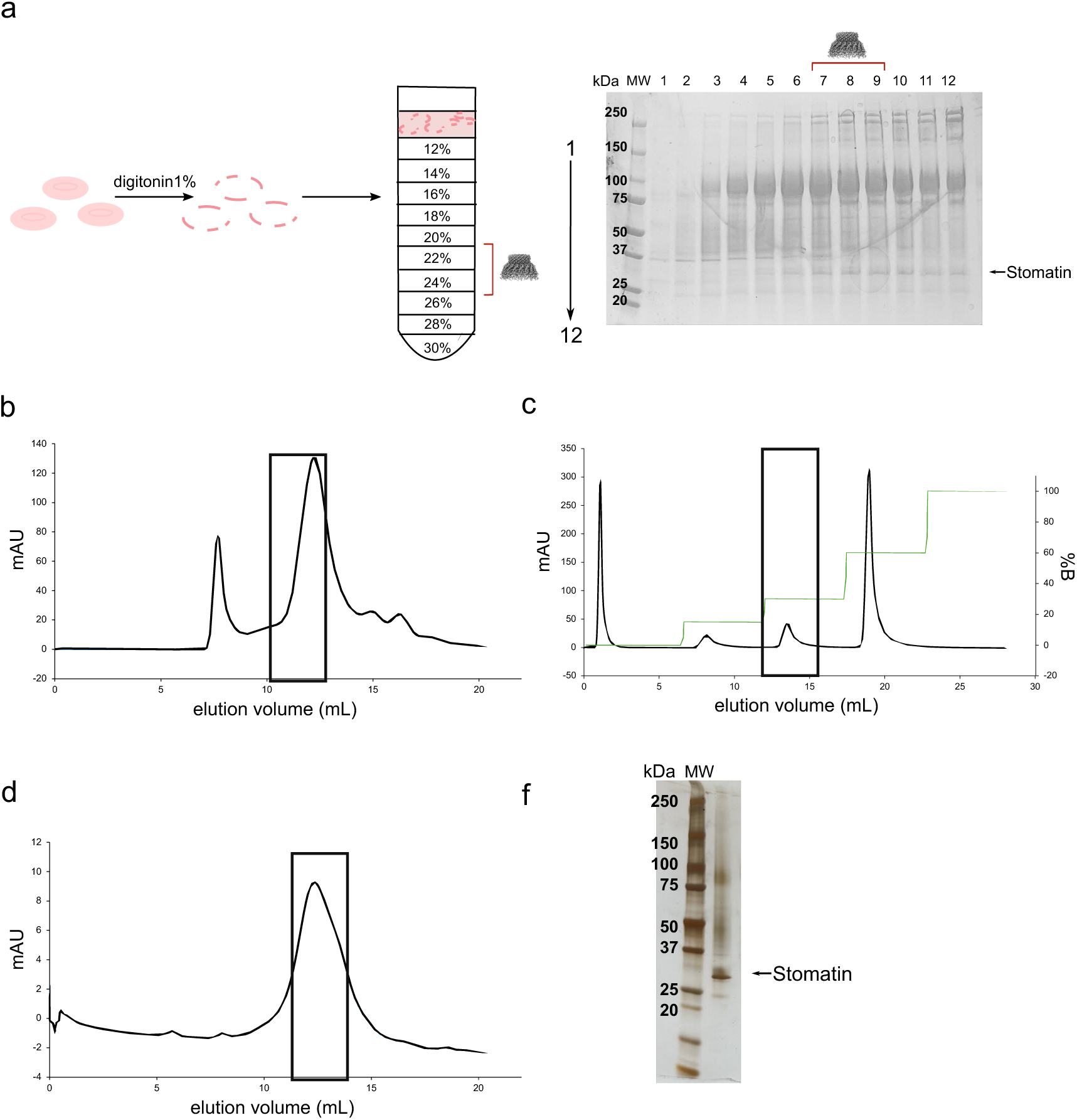
Purification of human stomatin. **a,** Human erythrocyte ghost membranes (pink empty erythrocytes) were washed and solubilized with 1% (w/v) digitonin. Solubilized membranes were separated on a 12–30% glycerol gradient, and the collected fractions were analyzed by SDS–PAGE. A red bracket indicates fractions containing stomatin. **b,** Size-exclusion chromatography (Superose 6) of stomatin-containing fractions. The region enriched in stomatin (black box) was collected. **c** Collected fractions were subjected to ion-exchange chromatography with a HiTrap ANX FF column and eluted using 3 different gradients of KCl (150, 300, and 600 mM). **d,** Size-exclusion chromatography (Superose 6) of the fraction eluted with the 300 mM KCl (black box). **e,** SDS–PAGE gel, silver-stained, of the fraction corresponding to the peak from panel d.

**Extended Data Fig. 2:**
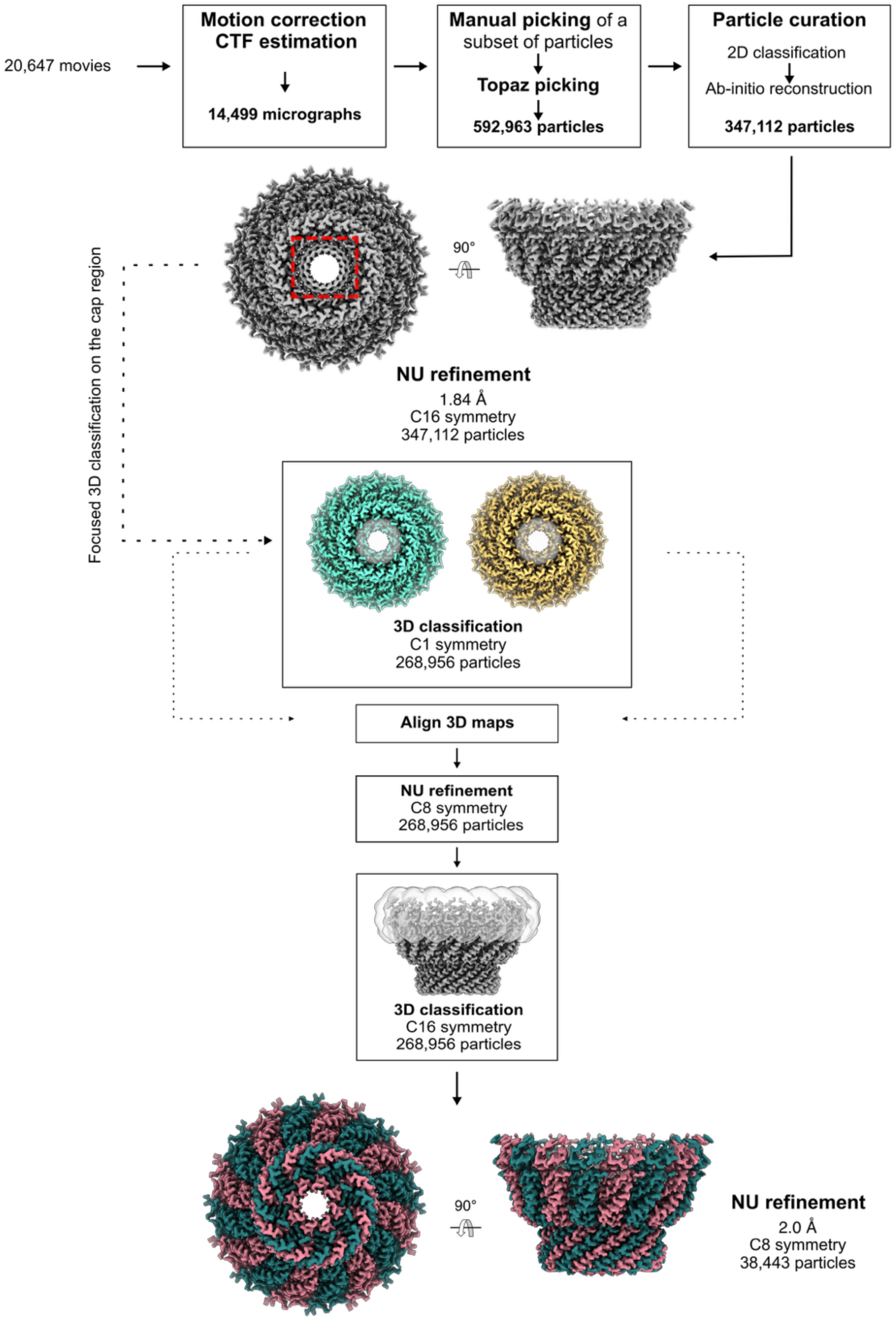
Cryo-EM workflow. Flowchart outlining cryo-EM image acquisition and processing performed to obtain the structure of stomatin. All processing was performed using CryoSPARC v4.4-v4.71 (see Methods for details).

**Extended Data Fig. 3:**
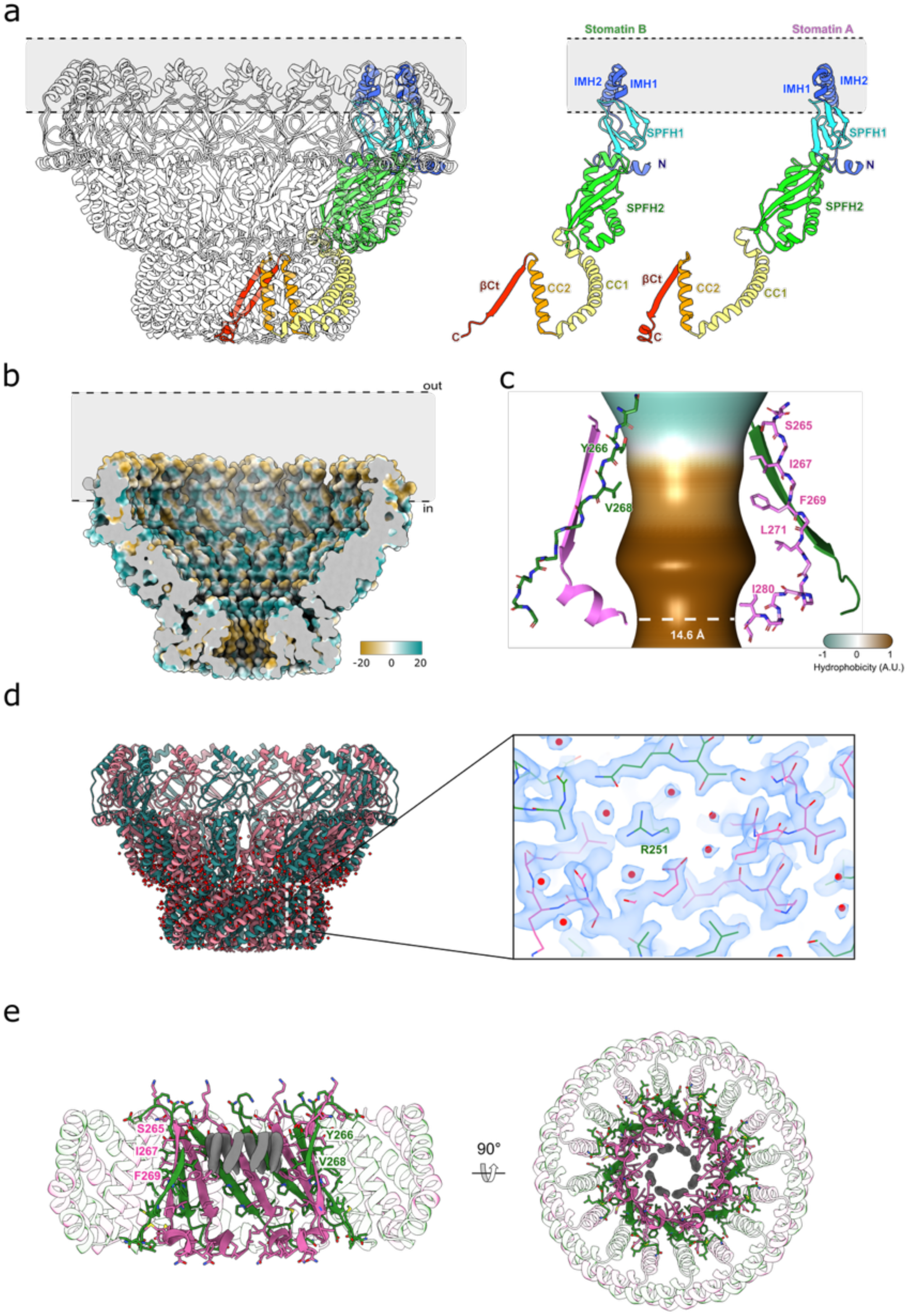
**a,** Stomatin oligomer showing a stomatin dimer colored within the assembly, with other protomers semi-transparent (left); the isolated protomers (right) outlines the stomatin topology. **b,** Cut-through view of the stomatin barrel in the membrane plane, surface-colored by lipophilicity potential (orange, hydrophobic; aquamarine, hydrophilic) as defined in ChimeraX, revealing the distribution of hydrophobic surfaces within the membrane-embedded region and the apical pore. **c,** Interior aperture of stomatin cap. The pore size and hydrophobicity of the barrel were calculated using the software CHAP^45^. **d,** Side (left) view of stomatin model, with assigned water molecules shown in red. A close up of the cryoEM density map with atomic model around R251 is shown, showing a hydrogen-bonded network of waters between the CC2 domain and the C-terminal beta barrel **e,** Close-up of the top pore from the side (left) and top (right), revealing eight unidentified lipidic densities (dark grey). Residues in close contact with these densities are labeled.

**Extended Data Fig. 4:**
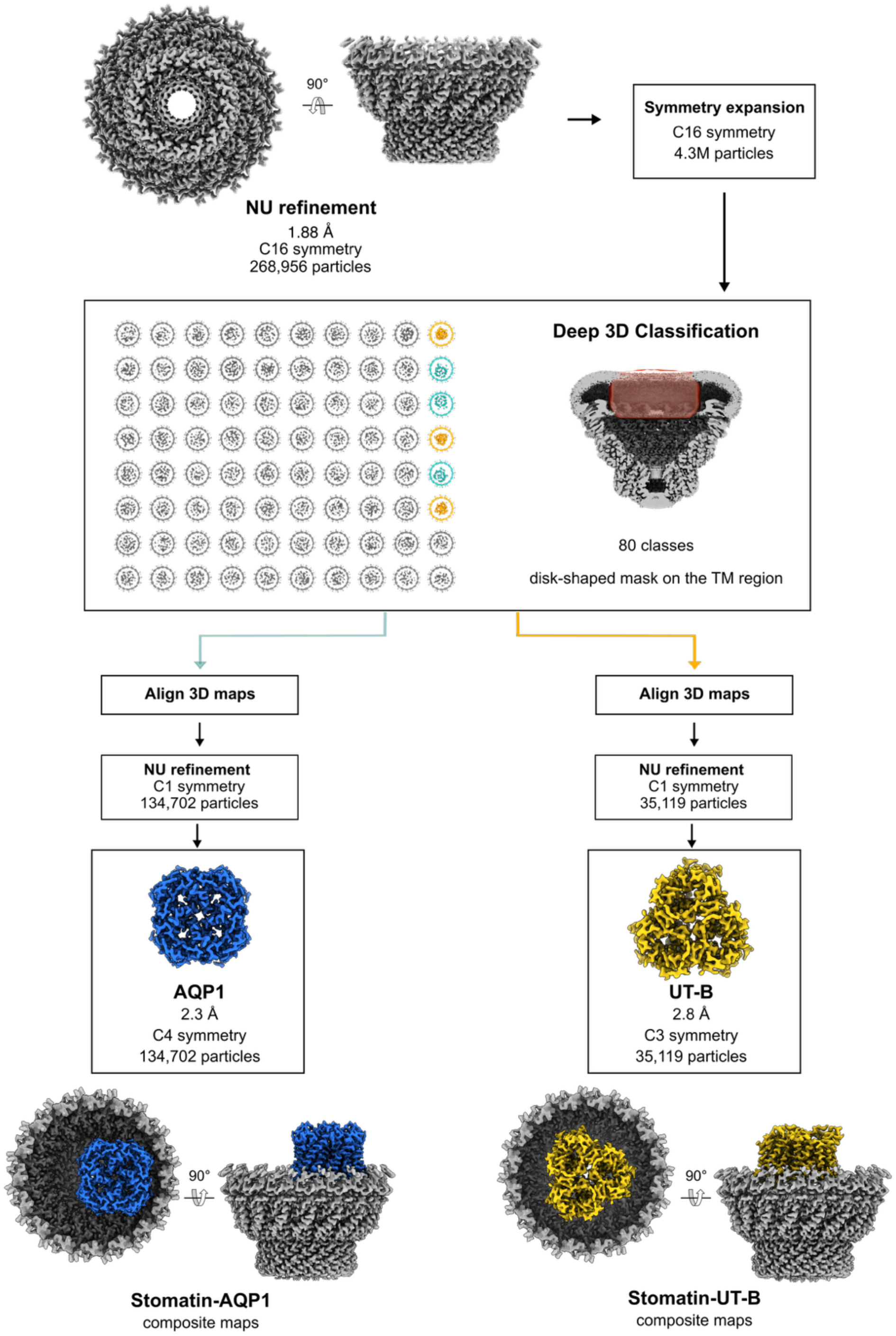
Cryo-EM workflow (AQP1 & UT-B complexes). Flowchart outlining cryo-EM image acquisition and processing performed to obtain the structure of endogenous stomatin complexes with AQP1 and UT-B. All processing was performed using CryoSPARC v4.4-4.71 (see Methods for details).

**Extended Data Fig. 5:**
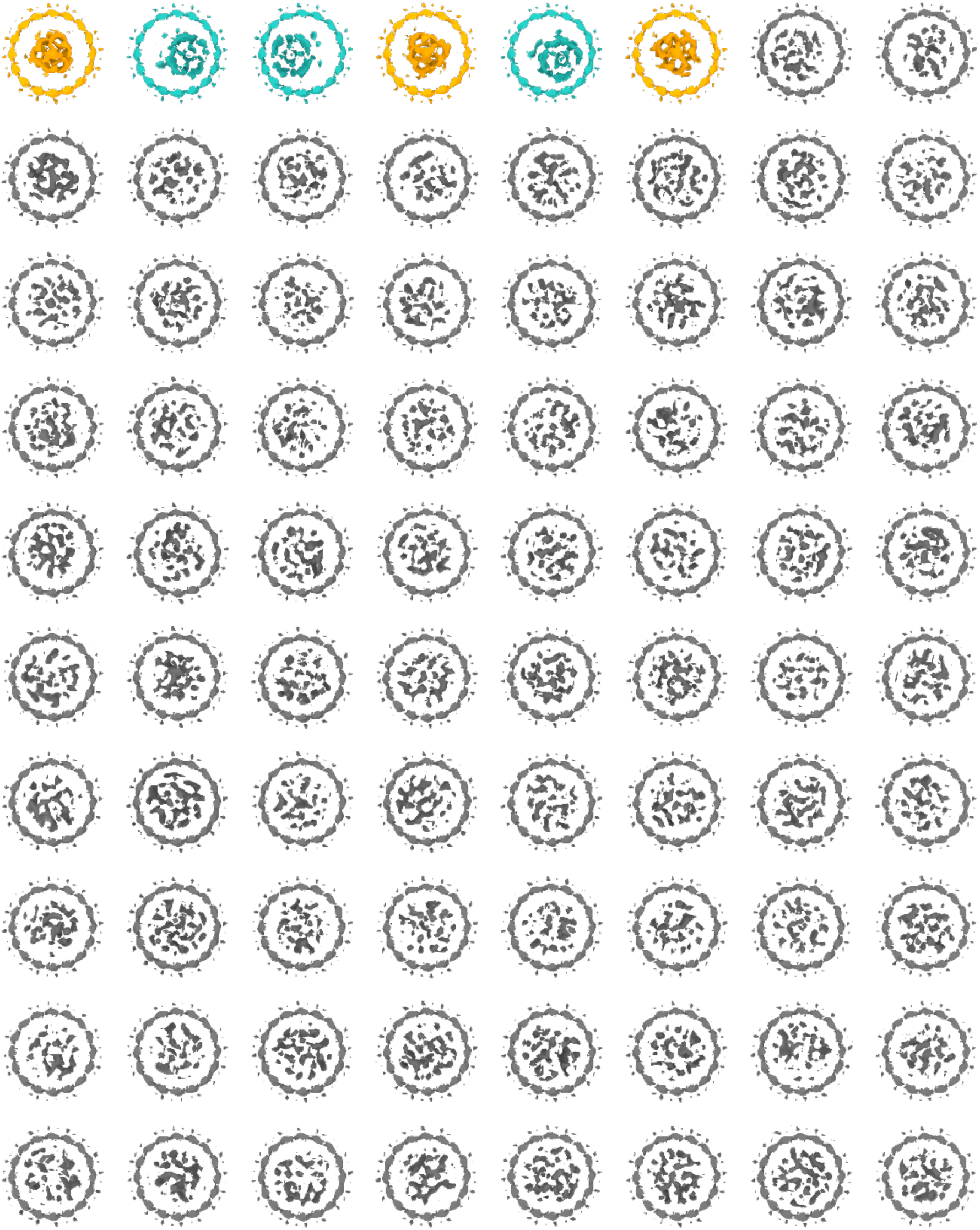
Deep 3D classification of stomatin complexes. Complete set of 80 classes shown with a cut-through view of the transmembrane region, obtained after 3D classification using a disk-shaped mask centered inside the stomatin intramembrane belt. Low-resolution features reveal two identifiable binding partners: a tetrameric complex consistent with AQP1 (light blue classes) and a trimeric complex corresponding to UT-B (yellow classes). Grey classes have uninterpretable density within the transmembrane region.

**Extended Data Fig. 6:**
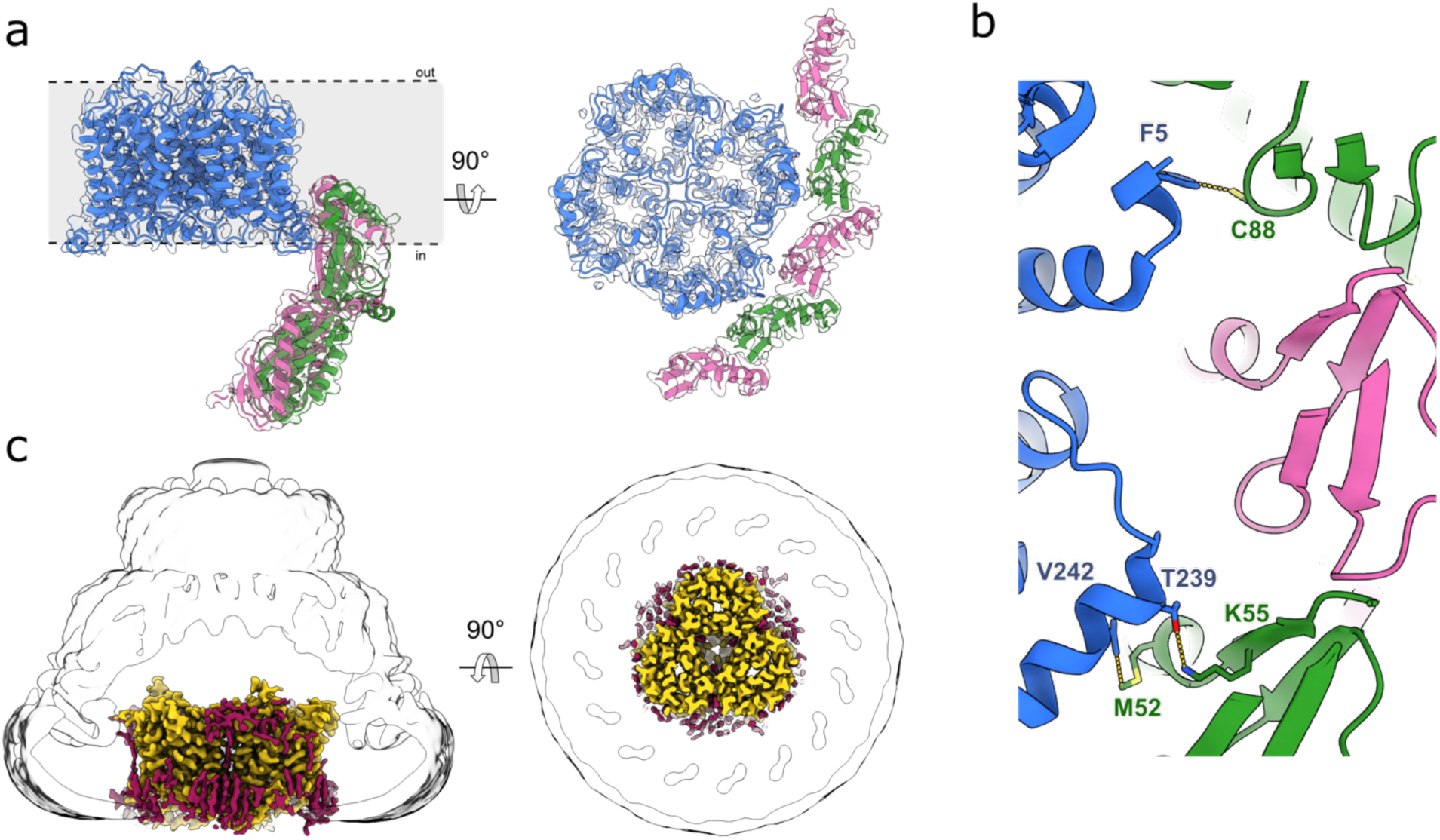
Interactions of stomatin with AQP1 and UT-B. **a**, Side (left) and cytosolic (right) views of AQP1 (blue) map showing how AQP1 interacts with stomatin. **b**, Close up of the interactions between stomatin (green and pink) and AQP1, showing key residues that are involved in the interaction. **c**, Side (left) and top (right) views of UT-B (yellow) map surrounded by lipids (dark red). The Gaussian-filtered delineating the micelle and stomatin molecular envelope is shown as a transparent white surface.

**Extended Data Fig. 7:**
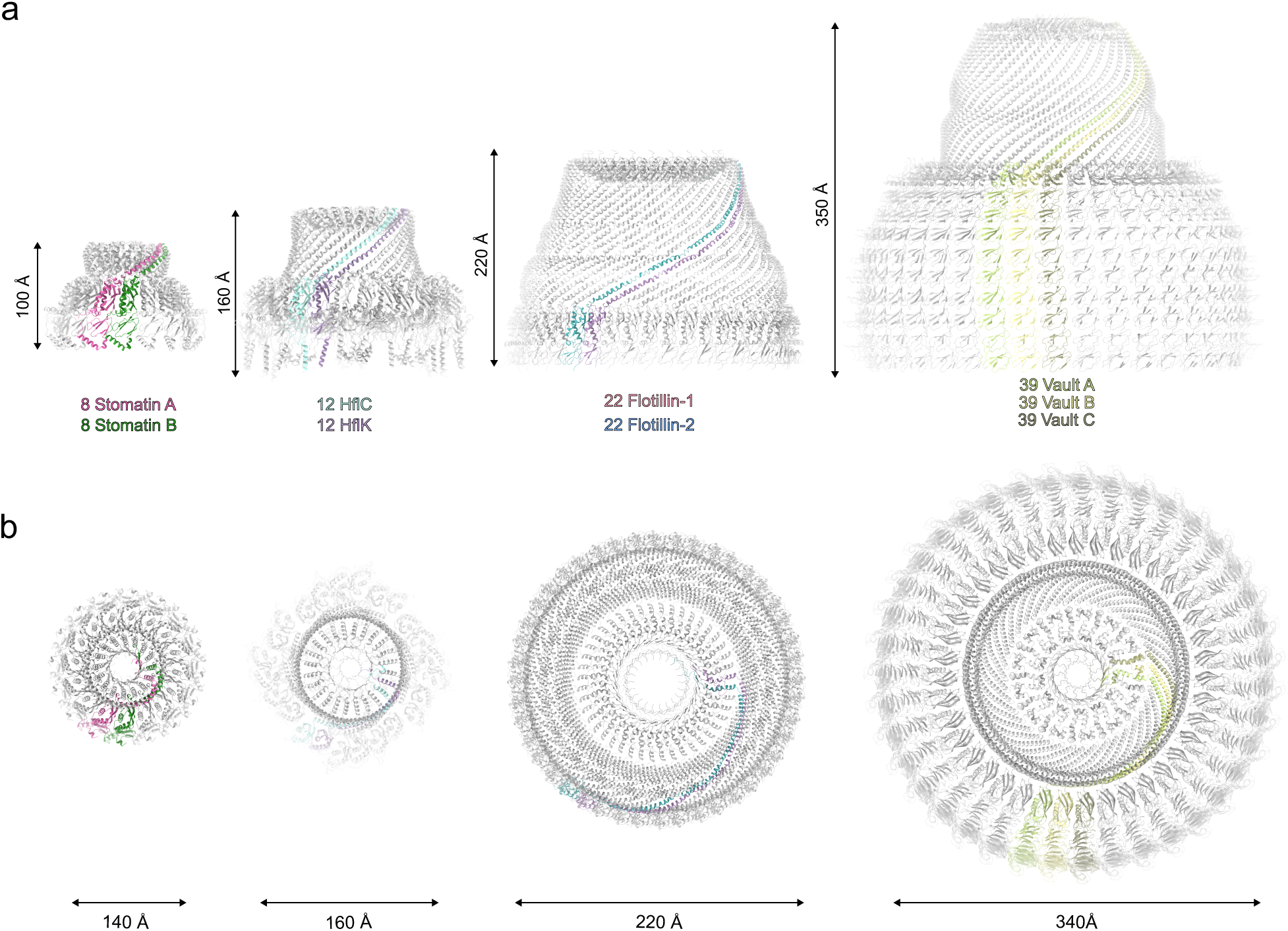
Comparison with other SPFH family members. **a,** Structures viewed in the membrane plane: stomatin (pink and green) compared with HflK/C (PDB: 7WI3), with HflK and HflC in two colors; flotillin (PDB: 9BQ2), with flotillin-1 and flotillin-2 in two colors; and a half-shell of major vault protein (PDB: TBD). **b,** Same structures as in panel a, viewed from the cytoplasmic side (or end-on, for vault).

**Extended Data Fig. 8:**
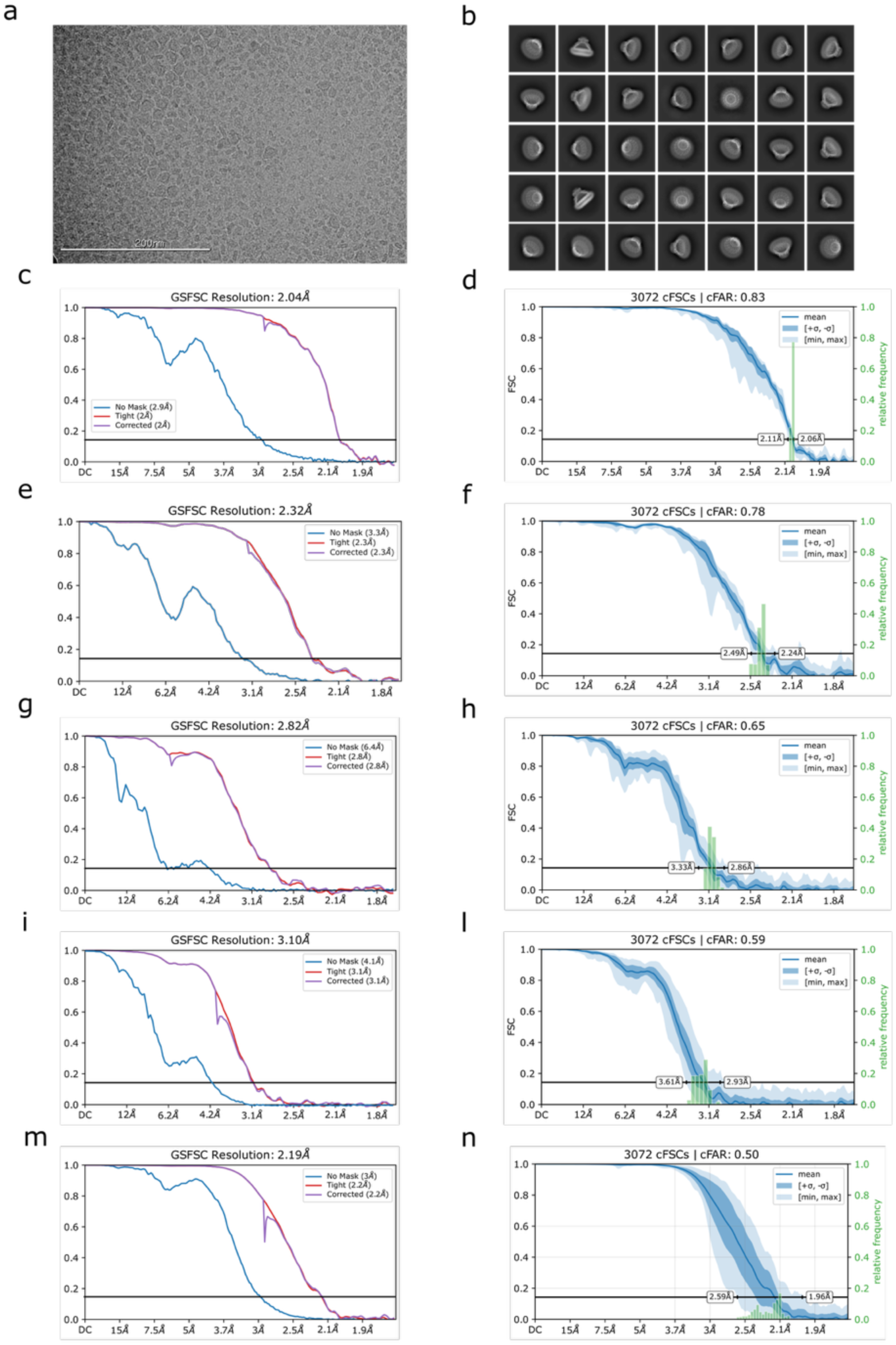
Single particle cryo-EM structure determination. **a,** Representative cryo-EM micrograph. **b,** 2D class averages of picked particles, ordered by population. **c,** Fourier shell correlation (FSC) curve for stomatin (C8 symmetry). **d,** Directional anisotropy of stomatin with C8 symmetry, calculated using Orientation Diagnostics in CryoSPARC 4.7.1. **e,** FSC curve for AQP1 local refinement in C4 symmetry. **f,** Directional anisotropy of the AQP1, calculated using Orientation Diagnostics in CryoSPARC 4.7.1. **g,** FSC curve for UT-B local refinement in C3 symmetry. **h,** Directional anisotropy of UT-B, calculated using Orientation Diagnostics in CryoSPARC 4.7.1. **i,** FSC curve for the local refinement of AQP1-stomatin interface in C1 symmetry. **l,** Directional anisotropy of AQP1-stomatin interface, calculated using Orientation Diagnostics in CryoSPARC 4.7.1. **m,** FSC curve for the local refinement of stomatin intramembrane in C1 symmetry. **n,** Directional anisotropy of stomatin intramembrane in C1 symmetry, calculated using Orientation Diagnostics in CryoSPARC 4.7.1.

**Extended Data Fig. 9:**
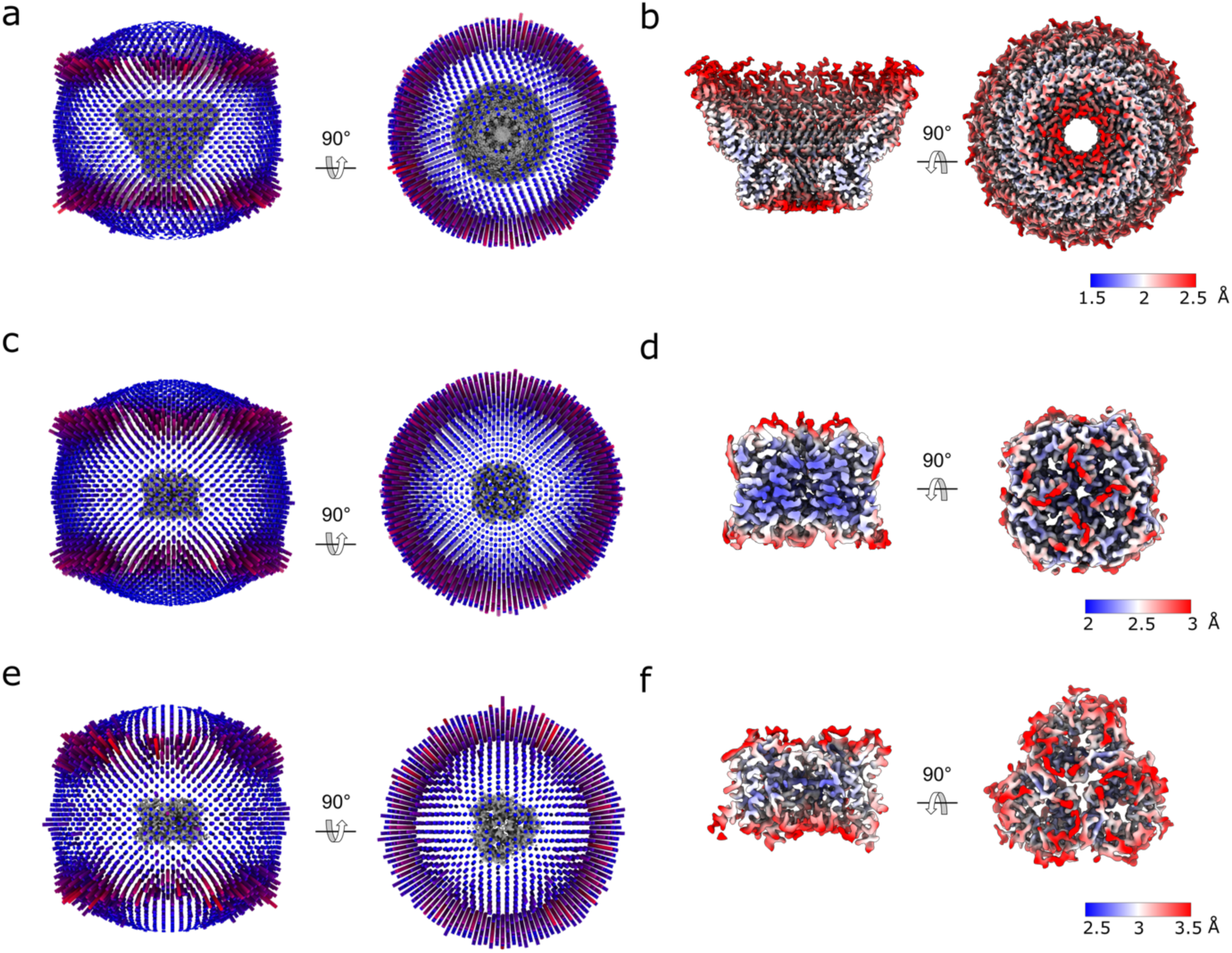
**a,** Orientation distribution plot for the 38k particles used in the stomatin C8 refinement. **b,** Local resolution plot of the cryoEM map from the global 3D reconstruction for stomatin. **c,** Orientation distribution plot for the 135k particles used in the AQP1 refinement visualized using `star2bild.py` from `pyem`^46^. **d,** Local resolution plot of the cryoEM map from the global 3D reconstruction for AQP1. **e,** Orientation distribution plot for the 35k particles used in the UT-B refinement. **f,** Local resolution plot of the cryoEM map from the global 3D reconstruction for UT-B.

**Extended Data Fig. 10:**
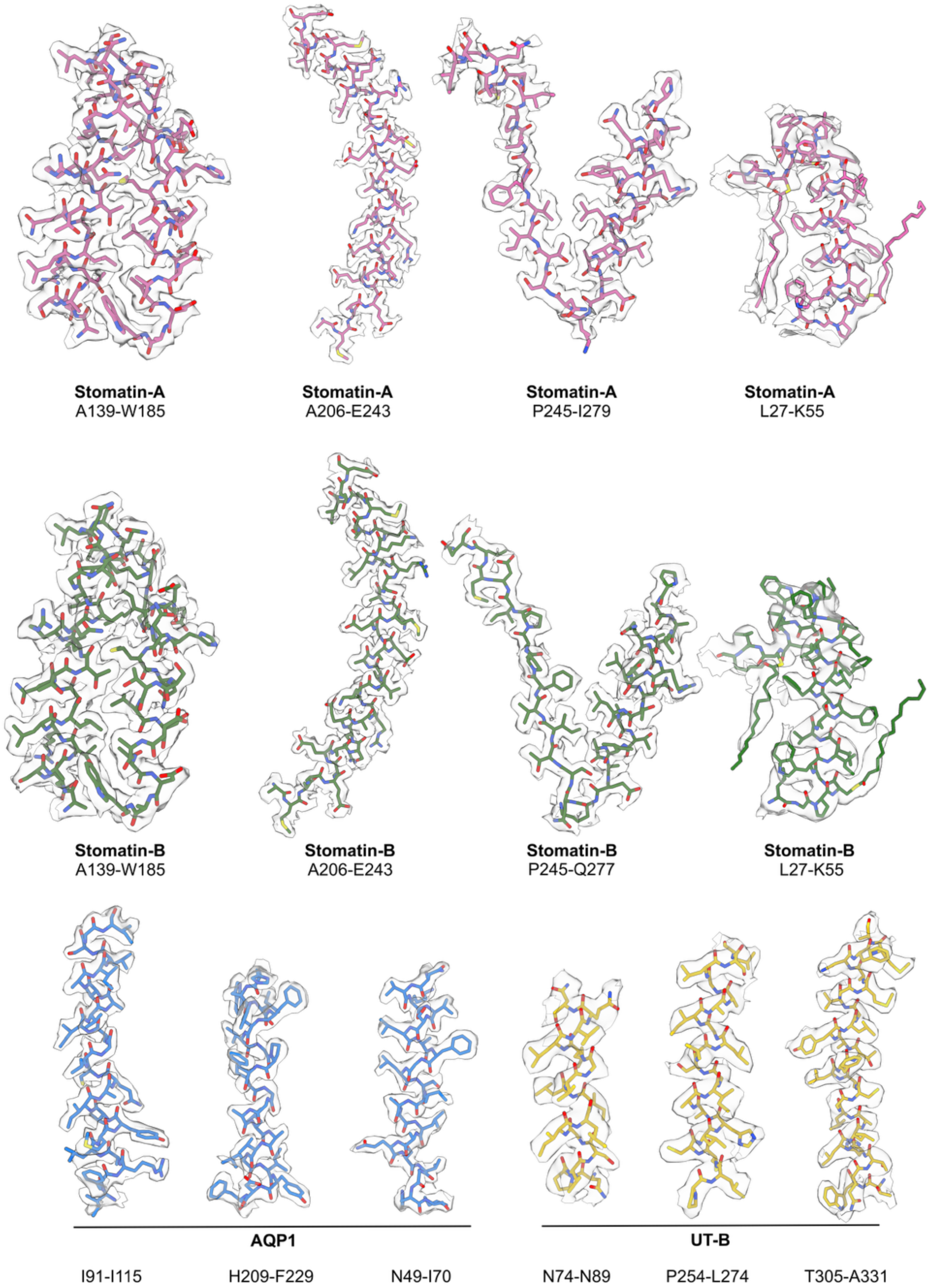
Model/map fit. Cryo-EM densities (transparent gray surface; generated using surface zone in ChimeraX) are shown with corresponding segments of the stomatin atomic model and for AQP1 and UTB; sidechains are rendered in stick representation and colored as in Figures 1 and 3. The first three panels for stomatin A & B are rendered using the sharpened map from the C8 refinement (EMD: TBD); The two showing the intramembrane helices are rendered using the unsharpened map from the local asymmetric refinement of SPFH1/IMH (EMD: TBD). The AQP1 model & map correspond to the sharpened map from the local C4 refinement (EMD:TBD) and the UT-B model and map correspond to the sharpened map from the C3 refinement (EMD:TBD).

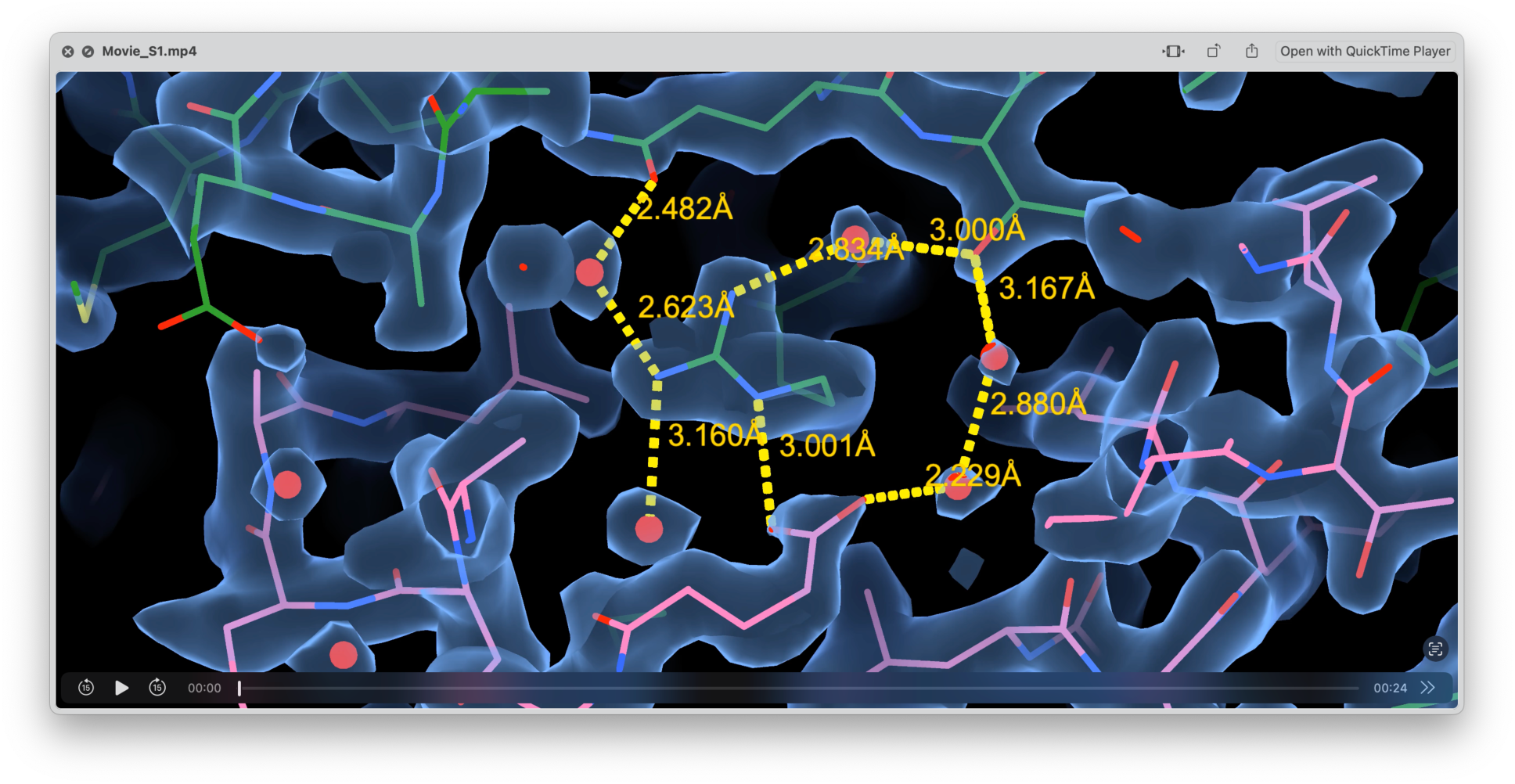

Movie S1: Hydrogen-bonded network of waters between the CC2 domain and the C-terminal beta barrel

Link

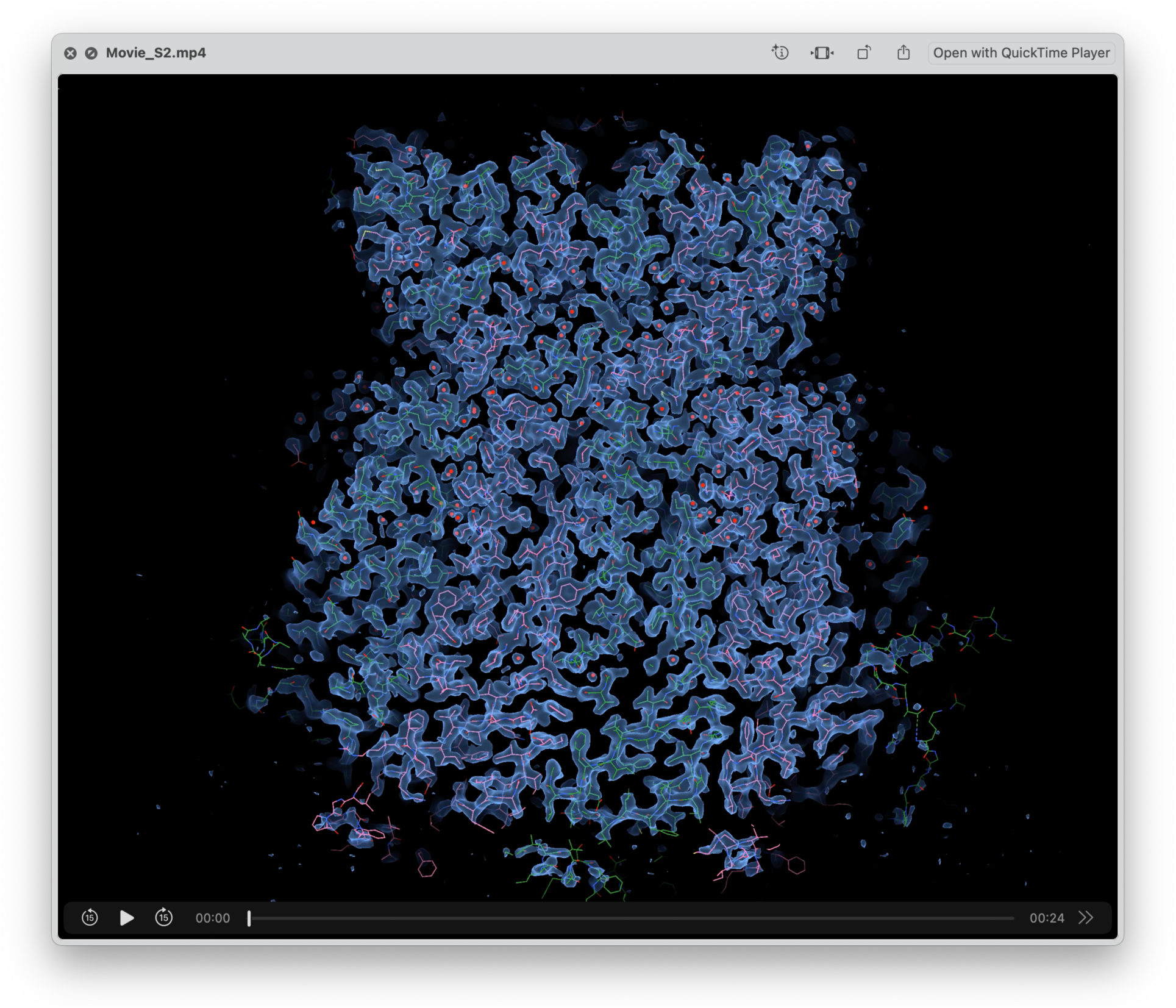

Movie S2: Overall map/model fit (C8 sharpened map)

Link

